# Comprehensive mapping of mammalian transcriptomes identifies conserved genes associated with different cell differentiation states

**DOI:** 10.1101/022608

**Authors:** Yang Yang, Yu-Cheng T. Yang, Jiapei Yuan, Xiaohua Shen, Zhi John Lu, Jingyi Jessica Li

## Abstract

Cell identity (or cell state) is established via gene expression programs, represented by “associated genes” with dynamic expression across cell identities. Here we integrate RNA-seq data from 40 tissues and cell types from human, chimpanzee, bonobo, and mouse to investigate the conservation and differentiation of cell states. We employ a statistical tool, “Transcriptome Overlap Measure” (TROM) to first identify cell-state-associated genes, both protein-coding and non-coding. Next, we use TROM to comprehensively map the cell states within each species and also between species based on the cell-state-associated genes. The within-species mapping measures which cell states are similar to each other, allowing us to construct a human cell differentiation tree that recovers both known and novel lineage relationships between cell states. Moreover, the between-species mapping summarizes the conservation of cell states across the four species. Based on these results, we identify conserved associated genes for different cell states and annotate their biological functions. Interestingly, we find that neural and testis tissues exhibit distinct evolutionary signatures in which neural tissues are much less enriched in conserved associated genes than testis. In addition, our mapping demonstrate that besides protein-coding genes, long non-coding RNAs serve well as associated genes to indicate cell states. We further infer the biological functions of those non-coding associated genes based on their co-expressed protein-coding associated genes. Overall, we provide a catalog of conserved and species-specific associated genes that identifies candidates for downstream experimental studies of the roles of these candidates in controlling cell identity.

**Highlights:**

- Comprehensive transcriptome mapping of cell states across four mammalian species
- Both protein-coding genes and long non-coding RNAs serve as good markers of cell identity
- Distinct evolutionary signatures of neural and testis tissues
- A catalog of conserved associated protein-coding genes and lncRNAs in different mammalian tissues and cell types

## Introduction

Cell differentiation states or cell identities (e.g., embryonic stem cells (ESC), heart tissues, and HeLa cell line) are maintained and controlled by a set of key regulators and epigenomic modifications (1-3). Previous studies have revealed crucial roles of some key regulators in controlling gene expression during cell differentiation and developmental processes, including transcription factors (TFs) (3), chromatin regulators (4, 5), splicing proteins (6, 7), microRNAs (8, 9), and long non-coding RNAs (lncRNAs) (10, 11). For instance, transcription factors Oct4, Sox2 and Nanog collaboratively activate protein-coding and microRNA genes in ESCs (12, 13); several microRNAs play important roles in controlling self-renewal in ESC or initiating differentiation (14, 15); two novel lncRNAs serve as important regulators of heart development in mouse (16, 17). In addition, epigenomic modifications are also important in regulating transcription. For example, H3K27ac-marked super-enhancers are associated with key TF genes that control cell states (18, 19).

In developmental biology and genomics, an important question is to understand how individual biological molecules and their interactions determine cell states. Although several regulatory circuits have been found evolutionarily conserved in mammals (20, 21), it remains challenging to systematically identify conserved protein-coding genes and lncRNAs that function in various cell states across multiple mammalian species. Given the vast transcriptomic data produced in recent years, comparative analysis of mammalian transcriptomes becomes feasible to identify conserved genes and so reveal molecular mechanisms underlying cell identity control. Early comparative studies of mammalian transcriptomes were mainly based on microarrays and thus restricted to annotated protein-coding genes in closely related mammals (22, 23). More recent high-throughput RNA sequencing (RNA-seq) studies, though, have been able to comprehensively characterize both protein-coding and long non-coding genes’ expression patterns in multiple species (24), providing new insights into the evolution of gene regulation (24-26). For example, it was found that unlike protein-coding transcripts, the number of non-coding transcripts increases consistently with the phenotypic complexity of species (27). This suggests that non-coding RNAs might play critical roles in the evolution of eukaryotes.

In this study, we aim to systematically identify evolutionarily conserved genes that define cell identities in multiple mammalian species, as well as to find possible correspondences between cell identities across different species. We collected and processed 307 publicly available polyadenylated RNA-seq data sets for 40 tissues and cell types (including ESCs, iPSCs, *in vivo* tissues, and cultured cell lines) from four mammalian species (human, chimpanzee, bonobo, and mouse). In order to quantitatively characterize the lineage relationships of those tissues and cell types, we used a statistical method TROM (Transcriptome Overlap Measure) (28, 29) to find their correspondence by a comprehensive transcriptome mapping. The mapping results enabled us to construct a cell differentiation tree, to reveals novel relationships among cell identities in addition to known relationships. The results also provide a catalog of protein-coding and long non-coding associated genes, which well capture transcriptome characteristics of various cell identities in different species. In our investigation of the results, we found that different cell identities exhibit different levels of conservation in terms of protein-coding gene expression. For example, testis tissues are more highly enriched with the conserved protein-coding associated genes than neural tissues. Moreover, our analyses revealed that the conserved protein-coding associated genes are highly enriched in biological functions and cellular pathways, which are closely related to the physiology of their associated cell states. There is also enrichment of master transcription factors, which have been reported to determine cell identity (18, 19), in these conserved protein-coding associated genes, Those results suggest that our identified conserved protein-coding associated genes are good markers of cell identities. In addition to protein-coding genes, we found that lncRNAs also serve well as associated genes for establishing a good correspondence of cell identities across species. We inferred the potential functions of those lncRNAs from the known biological functions of protein-coding genes by constructing an evolutionarily conserved gene co-expression network. Interestingly, we found that the conserved associated lncRNAs exhibit significant enrichment of biological functions related to cell identities, suggesting that these lncRNAs have conserved functions in evolution and are also good markers of cell identities. Our study demonstrates that, by integrating and re-analyzing large-scale public transcriptomic data from multiple species using proper statistical methods, we are able to systematically discover unknown markers of cell identities and provide insights into their molecular and evolutional characteristics.

## Results

### 1. Identification of associated genes for different cell states

We first collected and uniformly processed 307 publicly available polyA RNA-seq data sets (∼19 billion sequencing reads) of various tissues and cell types from four mammalian species (mouse and three primates including human, bonobo and chimpanzee) (Figure 1A and Supplementary Figure 1). Among them, the 183 human and 77 mouse data sets span a wide range of developmental stages and lineages. The chimpanzee and bonobo data sets include iPSC and several *in vivo* tissues (brain/cortex, cerebellum, heart, kidney, liver, and testis) (Supplementary File 1).

**Figure 1.**
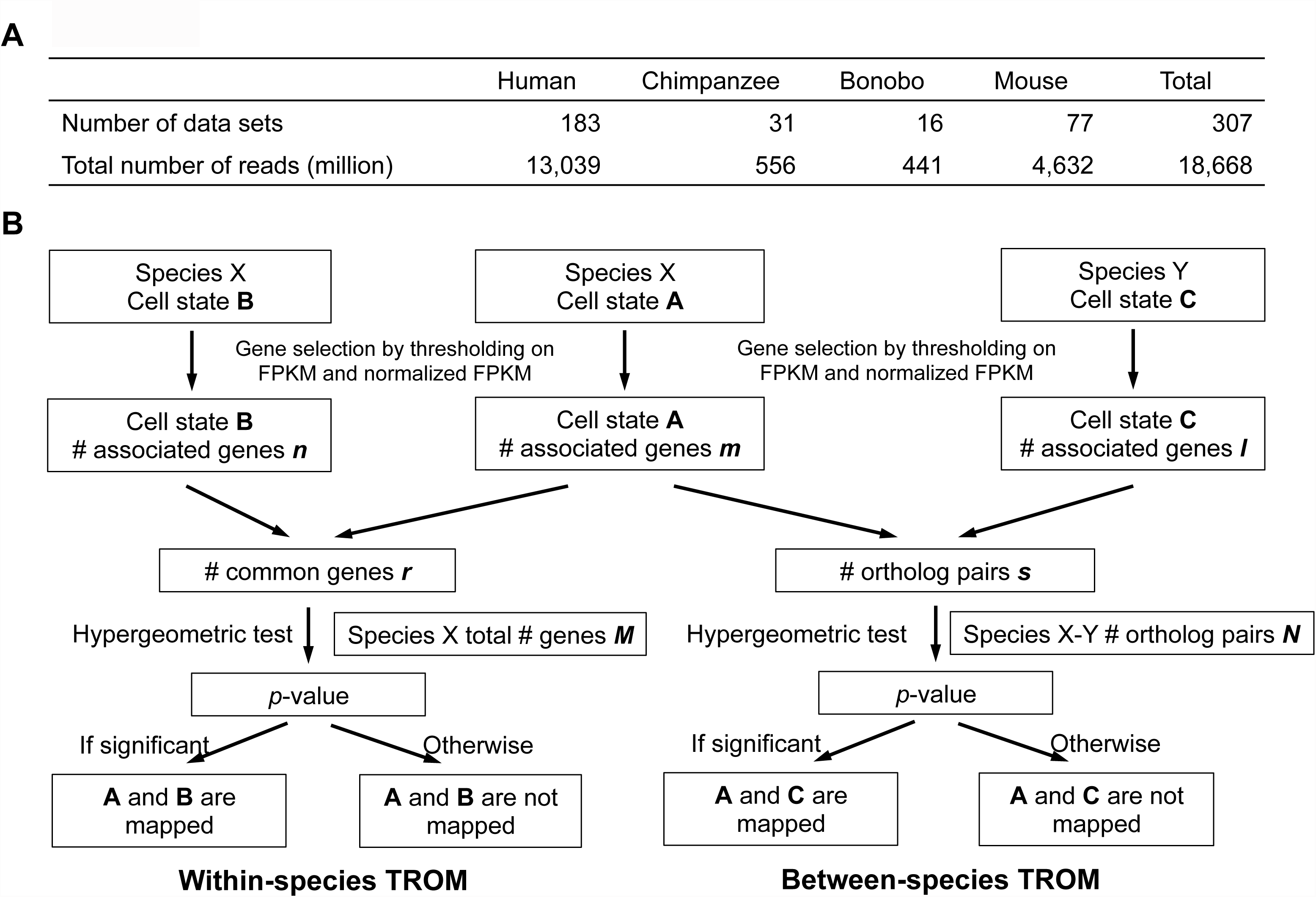
Overview of the RNA-seq data sets and TROM approach. (**A**) Numbers of RNA-seq data sets and sequencing reads for each mammalian species, including human, chimpanzee, bonobo, and mouse. (**B**) The TROM approach. First, associated genes of each cell state are selected using thresholds on FPKMs and *Z*-scores (normalized FPKMs across cell states). In the within-species TROM (left panel), the significance of the number of the common associated genes of two cell states is established via a hypergeometric test. In the between-species TROM (right panel), a similar hypergeometric test is carried out, except that orthologous genes are used to connect the two species. Two cell states are called “mapped” if the test is significant. (See Methods for details.)

To capture the transcriptome characteristics of different cell states, we define the “associated genes” of a cell state as the protein-coding and long non-coding genes whose expression is relatively high in that cell state and relatively low in some other cell states. To identify the associated genes of different cell states and subsequently use them to compare the cell states from different species, we adapted a statistical method TROM (Transcriptome Overlap Measure), which were recently developed for comparing developmental stages of *D. melanogaster* and *C. elegans* (28, 29). We first identified the associated genes of cell states by the following criterion: in a given cell state, its associated genes must have FPKM (fragments per kilobase of transcript per million mapped reads (30)) above a positive constant *c* (*c* = 1 for protein-coding genes and *c* = 0 for lncRNAs, because lncRNAs are generally more lowly expressed than protein-coding genes) and *Z*-score (normalized FPKM across samples) in the top 5% compared to its *Z*-scores in all cell states (see Methods for details, Supplementary Figure 2). We then used those identified associated genes to compare the cell states within and between mammalian species (Figure 1B). Specifically, for every pair of cell states, we tested if there is significant overlap between their associated genes. If so, the two cell states are called “mapped”. In a within-species comparison, the common associated genes shared by two cell states is considered the overlap; in between-species comparisons, the overlap is defined as the orthologous genes pairs between the associated genes of the two cell states, each from a different species (Supplementary File 2). In our study, protein-coding genes were used to compare all the four species, while lncRNAs were only used in the comparisons of the three primate species, because orthologous lncRNAs are largely unknown between primates and mouse. Like other comparative genomic studies, we also used correlation analyses to compare different cell states based on measured gene expression levels. However, in contrast to TROM, these correlation analyses failed to find clear or informative correspondence patterns among the mammalian cell states (Supplementary Figure 3). This was expected and consistent with our previous findings (28), because correlation values depend heavily on the accuracy of gene expression estimates. On the other hand, TROM finds correspondence between cell states based on their associated genes and is more robust to noise and biases in gene expression estimates.

## 2. Within-species cell state mapping

### 2.1 Lineage relationships rediscovered by cell state correspondence maps

We first applied TROM to map transcriptomes of human cell states (i.e., tissues and cell types) using their associated protein-coding genes, resulting in a clear correspondence map (Figure 2A). As expected, transcriptomes of the same or similar tissue types form prominent mapping blocks, including (i) ESCs and iPSCs, (ii) cultured cell lines, (iii) immune cells and tissues, (iv) neural tissues, (v) liver tissues, and (vi) testis tissues. These blocks are consistent with the known physiology of tissues and cells. For example, similar to ESCs, iPSCs are capable of generating all types of differentiated tissues and cells.

**Figure 2.**
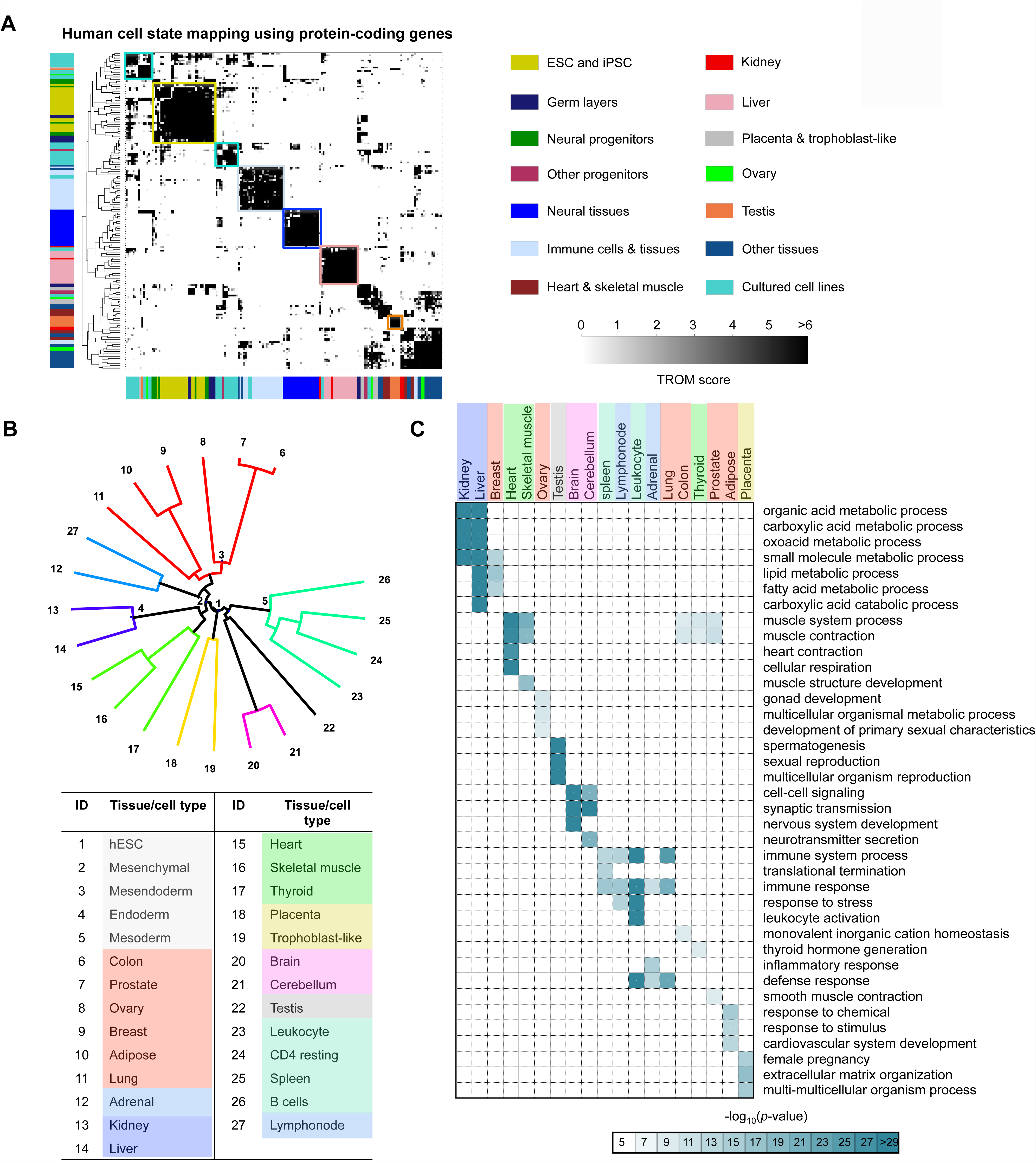
Cell states encoded by associated protein-coding genes in human. (**A**) A correspondence map of human cell states by TROM using associated protein-coding genes. Columns and rows correspond to biological samples of various cell states. More significant TROM scores (defined as –log_10_ transformed Bonferroni corrected *p*-values from hypergeometric tests) are shown in darker colors. Axis colors represent cell states, and colored boxes mark the prominent mapping patterns. (**B**) Cell differentiation tree constructed using associated protein-coding genes of 27 human tissues and cell types. Five progenitor cell types (ID 1-5) are placed in the origin by the Hungarian algorithm. Colors are used to distinguish cell state clusters. (**C**) Enriched GO (biological processes) terms of 19 human cell states. More significant enrichment scores (defined as –log_10_ transformed *p*-values) are shown in darker colors. ell states are marked with the same colors as in (B).

Next we compared those cell states again by TROM using only associated TFs or associated lncRNAs (Supplementary Figure 4). Interestingly, the resulting mapping patterns are similar to Figure 2A but have smaller mapping blocks, confirming the higher tissue specificity of TFs (31) and lncRNAs (32) than protein-coding genes. For example, placenta, trophoblast-like cells, and mesenchymal stem cells form a clear mapping block when compared using TFs. This observation can be explained by the fact that trophoblasts develop into a large part of the placenta, and placenta-derived cells have mesenchymal stem cell potentials (33). These results suggest that the identified associated TFs and lncRNAs serve well as representatives of cell identity, consistent with previous findings on their context specific activations and regulatory roles (31, 32).

In addition to human, we compared the transcriptomes of cell states within other mammalian species using all the associated protein-coding genes, only the associated TFs or the associated lncRNAs (Supplementary Figure 5-7). The mapping patterns are largely consistent with what we observed in human. Those within-species mapping results demonstrate that our approach is powerful and robust in revealing cell state similarity in multiple mammalian species, and justify our identified associated genes as good markers of cell identity.

### 2.2 Cell differentiation tree constructed using cell state associated genes

A cell differentiation tree is a good representation of the developmental genealogy of cell states. However, an accurate cell differetiation tree has not previously been constucted using RNA-seq data. Here, based on our identified associated genes, we constructed a tree of human cell states using a similarity metric defined as the percent of common associated genes between two cell states in the union of their associated protein-coding genes) (see Methods for details). We found that the resulting tree reflects well the known hierarchical lineage relationships of the human tissues and cell types (Figure 2B). Compared to terminally differentiated tissues, embryonic germ layers (ectoderm, mesoderm, endoderm, and mesendoderm) are more proximal to the ESC and pluripotent stem cells (e.g., mesenchymal stem cells). Moreover, the various terminally differentiated tissues exhibit biologically meaningful groups in the tree: (i) liver and kidney (blue), (ii) placenta and trophoblast-like cells (yellow), (iii) heart, skeletal muscle, and thyroid (emerald), (iv) immune cells and tissues (green and cerulean), (v) neural tissues (pink), (vi) testis (black), and (vii) other tissues (red).

Since TFs are considered more tissue-specific than non-TF protein-coding genes, we constructed this tree again using only the associated TFs. The resulting TF-based differentiation tree (Supplementary Figure 8) is similar to the previous tree in Figure 2B, with a few small changes. For example, lymphonode is now clustered with other immune cells and tissues, while it was grouped with adrenal in Figure 2B. Testis now forms an isolated terminally differentiated lineage, while it was in the same lineage as brain and cerebellum. This indicates the distinct gene expression pattern of TFs in testis, consistent with the literature (25). Since our constructed cell differentiation trees are reasonable and agree with prior knowledge on lineage relationships, they further justify our identified associated genes as good cell state markers.

### 2.3 Biological functions of cell state associated protein-coding genes

To investigate the biological functions of our identified associated genes, we started with the associated protein-coding genes, which have more complete functional annotations than lncRNAs. We first assayed the enriched Gene Ontology (GO) terms of the associated protein-coding genes for various cell states in human (Figure 2C) and mouse (Supplementary Figure 9). The results revealed that the associated protein-coding genes are enriched with biological processes that largely define the identities of the respective cell states. For example, kidney and liver are enriched with metabolic processes (including organic acid metabolic process, carboxylic acid metabolic process, oxoacid metabolic process and small molecule metabolic process), and testis is enriched with spermatogenesis and sexual reproduction. Interestingly, although kidney and liver share some metabolic processes, liver is additionally enriched with processes related to lipid and fatty acid metabolism. In addition, we found a great similarity between those GO terms enriched in associated genes and the “tissue-specifically enriched GO terms” identified using super-enhancer-associated genes in another study (18). For example, GO terms such as “muscle contraction”, “muscle system process” and “heart contraction” are enriched in the associated genes of heart and skeletal muscle in our study and the previous study (18). This observation suggests that our identified associated genes have a strong association with the super-enhancers and possibly other *cis*-regulatory elements.

We next identified the enriched KEGG cellular pathways enriched in the associated protein-coding genes of different cell states in human (Supplementary Figure 10) and mouse (Supplementary Figure 11). The resulting enriched cellular pathways are biologically meaningful. Among them an interesting result is that heart-associated genes are highly enriched with pathways related to neurodegenerative diseases (e.g., Alzheimer’s disease, Parkinson’s disease, and Huntington’s disease). The reason is that a significant number of heart-associated genes encode NADH dehydrogenase subunits and Cytochrome C oxidase subunits, which are involved in Alzheimer’s disease by previous biochemical evidence (34). This observation is also supported by previous clinical studies, which suggest that heart failure is linked to the development of neurodegenerative diseases (35).

Given those identified enriched GO terms and cellular pathways, we further characterized their dynamics during the differentiation process from stem cells to differentiated neural cells. We selected the following tissues and cell types to represent this differentiation progress: (i) ESC/iPSC, (ii) embryonic germ layers (including mesendoderm, ectoderm, mesoderm, and endoderm), (iii) neuroectodermal spheres and neural progenitor cells, (iv) differentiated neural tissues (including brain and cerebellum). By investigating the enriched biological functions at each of those cell states, we observed a clear shift in biological processes as the differentiation goes on (Supplementary Figure 12), consistent with previous studies (36). For example, cell states closer to the stem cell end, such as the embryonic germ layers, are enriched with basic metabolic processes; on the other hand, fully differentiated neural tissues contain enriched functions on neuronal development and signaling.

## 3. Between-species cell state mapping

### 3.1 Conserved expression patterns revealed by between-species cell state correspondence

As the within-species results have shown the effectiveness of our approach in mapping cell states and finding cell state markers, we further extended this approach to comparing cell states in different mammalian species. We evaluated the similarity of two cell states from different species by adopting TROM, which is based on the number of shared orthologous genes in the associated genes of the two cell states (see Methods for details). We first performed a comprehensive mapping between human and mouse cell states based on their associated protein-coding genes (Figure 3A). Several prominent mapping blocks emerged between corresponding tissues in human and mouse, including (i) neural tissues, (ii) heart and skeletal muscle tissues, and (iii) testis tissues.

**Figure 3.**
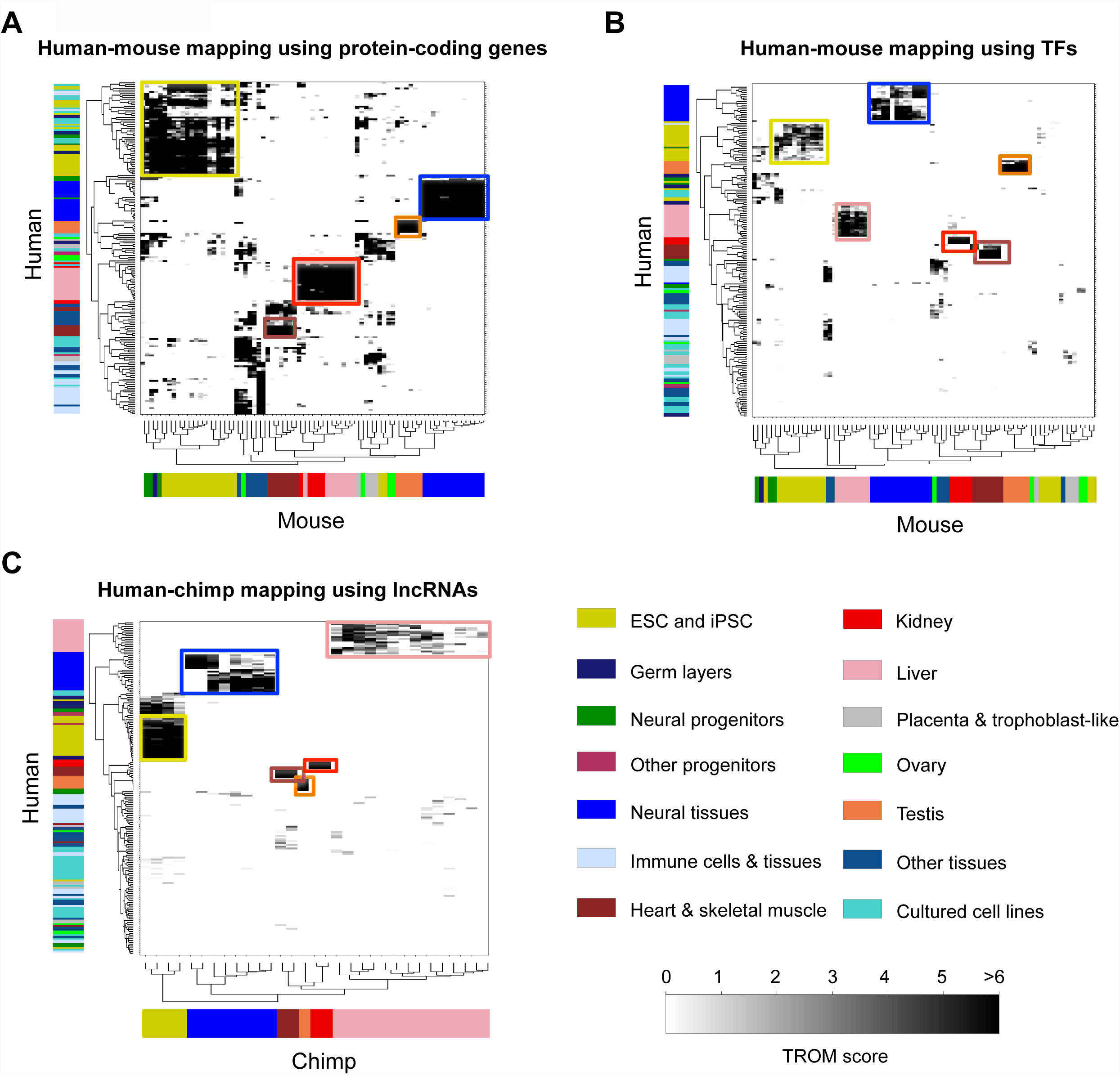
Cell state correspondence maps between human and other mammalian species. (**A**) A correspondence map of various cell states between human and mouse by TROM using associated protein-coding genes. Rows correspond to human cell states, and columns correspond to mouse cell states. (**B**) A correspondence map of various cell states between human and mouse by TROM using associated TFs. Rows correspond to human cell states, and columns correspond to mouse cell states. (**C**) A correspondence map of various cell states between human and chimpanzee by TROM using associated lncRNAs. Rows correspond to human cell states, and columns correspond to chimpanzee cell states. In (A)-(C), more significant TROM scores (defined as –log_10_ transformed Bonferroni corrected *p*-values) are shown in darker colors. Axis colors represent cell states, and colored boxes mark the prominent mapping patterns.

In addition to those expected mapping patterns, interesting mappings between tissues and cells of different types are also observed. The two most prominent mapping patterns are (i) human ESCs, iPSCs and cancer cell lines vs. mouse ESCs, iPSCs, germ layers and neural progenitor cells (NPCs) and (ii) human liver and kidney tissues vs. mouse liver and kidney tissues. Such mapping patterns are consistent with the known physiology of those tissues and cell lines. For example, ESCs and cultured cancer cell lines are both characterized by rapid cell proliferation, short cell cycles and blocks in differentiation (37, 38), which can explain our observed transcriptome similarity between human ESCs, iPSCs and cancer cell lines and mouse ESCs and iPSCs. Our observed similarity between human and mouse liver and kidney transcriptomes is a new discovery. It is a reasonable result because high similarity has been reported between liver and kidney in terms of their conserved DNase I-hypersensitive sites between human and mouse genomes (39). These mapping patterns show that that orthologous genes are conserved not only at the sequence level but also at the transcription level in terms of gene expression patterns (40, 41). They also confirm that the associated protein-coding genes are good cell state indicators.

We next performed a similar human-mouse cell state mapping using only the associated TFs (Figure 3B). Similar to the within-species results, the mapping patterns obtained using TFs are more sparse with smaller mapping blocks than their counterparts observed using protein-coding genes. Different mapping patterns have emerged. For example, almost no mapping is found between human cultured cancer cell lines and mouse ESCs/iPSCs (yellow box in Figure 3B). This result is also in line with our previous observations in the within-species mapping using TFs, indicating that ESCs/iPSCs are distinct from cancer cell lines in terms of TF programs. Since lncRNAs are not well conserved between human and mouse, no mapping was found between human and mouse cell states using the associated lncRNAs (Supplementary Figure 15B).

We also compared the cell states of human and those of bonobo and chimpanzee using the associated protein-coding genes, TFs or lncRNAs (Figure 3C and Supplementary Figures 13-15). Since many lncRNAs are conserved between primates, using the associated lncRNAs lead to clear mapping patterns, which are highly similar to the mapping results based on the associated TFs. In those mapping results, tissues of the same type are mapped across primate species. A possible explanation of those observations is that gene expression patterns evolve at comparable rates in different mammalian species (25).

### 3.2 Evolutionary signatures of cell state associated genes

To study the evolutionary properties of the cell state associated genes, we identified conserved associated protein-coding genes of several cell states, including ESC/iPSC, brain, cerebellum, heart, liver, kidney and testis, across the three primates and mouse. The percentages of human cell state associated genes having orthologs in chimpanzee, bonobo, and mouse are illustrated in Figure 4A. We found that among the associated protein-coding genes of each cell state, the conserved genes are significantly enriched with known tissue-specific genes from the TiGER database (42). (Supplementary Figure 16). Note that our cell state associated genes are not restricted to the specific genes of that cell state; they may include the genes that are also associated with other cell states. Hence ideally the TiGER tissue-specific genes should be contained in the associated genes of the corresponding tissues. However, we found that small portions of the TiGER tissue-specific genes were not identified as the associated genes. Our detailed analysis revealed that those non-associated tissue-specific genes in fact had much lower expression in that tissue than in some other tissues or cell types (Supplementary Figure 17). A possible explanation is that the TiGER tissue-specific genes were identified using fewer tissue types than in our study, suggesting that our results may be useful for refining the TiGER databse.

**Figure 4.**
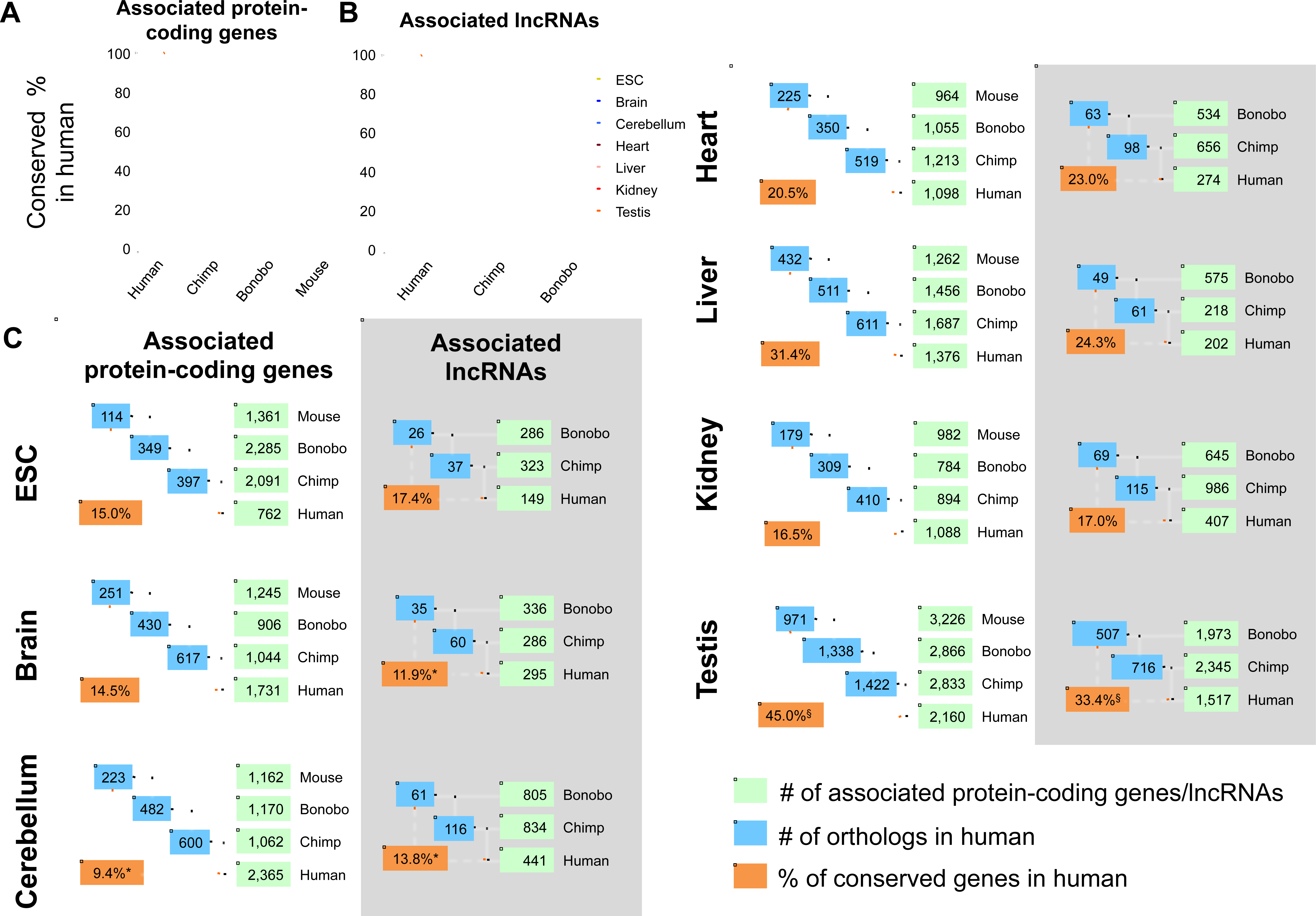
Conservation of associated protein-coding genes and lncRNAs in seven cell states. (**A**) Conservation percentages of human associated protein-coding genes of seven cell states (ESC, brain, cerebellum, heart, liver, kidney, and testis) in chimpanzee, bonobo and mouse. (**B**) Conservation percentages of human associated lncRNAs of seven cell states in chimpanzee and bonobo. (**C**) Sequential counts of human associated protein-coding genes and lncRNAs conserved in other species, from the closest to the most distant. Green boxes, numbers of all associated protein-coding genes and lncRNAs in each species. Blue boxes, counts of conserved associated protein-coding genes and lncRNAs. Orange boxes, percentages of conserved associated protein-coding genes and lncRNAs among all the human associated genes. The percentages, which are significantly higher or lower than expected (*p*-values<0.001, Fisher’s exact test), are marked with ^§^ or * respectively.

We next performed a sequential counting of the human associated genes conserved in other species, from the closest chimpanzee to the most distant mouse. This analysis is to reveal the conservation levels of different cell states based on our identified associated genes. We first observed interesting results on the protein-coding genes (Figure 4C). Among all the tissues and cell types, cerebellum has the smallest portion (9.4%; *p*-value<2.2e-16 by Fisher’s exact test) of associated protein-coding genes conserved in all the four species. This observation agrees with a previous finding that the evolution of protein-coding genes is more rapid in cerebellum than in other tissues (43). In contrast, testis contains the largest proportion (45.0%; *p*-value<2.2e-16 by Fisher’s exact test) of conserved associated protein-coding genes, confirming a previous observation that testis shows conserved functions and pathways in human and mouse (44). Since few conserved lncRNAs are known between primates and mouse (Supplementary Figure 18), we restricted our sequential counting of the conserved associated lncRNAs to the three primate species (Figure 4B). Similar to the associated protein-coding genes, the associated lncRNAs exhibit less conservation for neural tissues (11.9% for brain, *p*-value 4.84e-06 by Fisher’s exact test; 13.8% for cerebellum, *p*-value 2.06e-06) and the greatest conservation for testis (33.4%; *p*-value<2.2e-16 by Fisher’s exact test) (Figure 4C). These results suggest that the associated genes identified by TROM provides a simple and effective way to quantify evolution rates of various tissues and cell types simultaneously in an unbiased manner.

### 3.3 Conserved cell state associated genes are potentially key regulators

Previous studies revealed that a remarkably small number of key TFs could mainly define tissue-specific gene expression programs (3, 19). They have identified several sets of key TFs for certain human cell states (18). For example, Oct4, Sox2 and Nanog are considered essential in maintaining the ESC state and regulating embryonic development (3, 45, 46). We identified conserved associated TFs for seven cell states (Table 1, Supplementary File 3), and reasoned that these TFs should be functionally important based on our above analyses and previous literature. Most of these TFs have been reported as essential in tissue development and physiology (see the references of related literatures in Supplementary File 4). For example, we have successfully identified known self-renewal and pluripotency factors such as Oct4 (Pou5f1), Nanog, Sall4, Jarid2 and Myc (37) in ESCs as conserved associated TFs (Table 1). A recent study found that, compared to the commonly used “OSKM factors” (Oct4, Sox2, Klf4 and Myc), the “SNEL factors” (Sall4, Nanog, Esrrb and Lin28) can more efficiently generate iPSCs (47). Sall4, Nanog and Lin28 are successfully identified as conserved associated TFs in our analysis. Additionally, our conserved associated TFs in heart tissues significantly overlap with known lineage reprogramming factors that directly aid the conversion of fibroblasts to cardiomyocytes in human and mouse (Supplementary File 5) (48). Out of the six common reprogramming factors (GATA4/Gata4, HAND2/Hand2, MEF2C/Mef2C, MESP1/Mesp1, MYOCD/Myocd and TBX5/Tbx5) for human/mouse cardiomyocytes, we identified GATA4/Gata4, HAND2/Hand2, MEF2C/Mef2C, MYOCD/Myocd and TBX5/Tbx5 as conserved associated TFs. More importantly, some of those conserved associated TFs have not been well studied and could serve as interesting candidates for further functional studies on reprogramming or lineage conversion.

**Table 1.** Conserved cell state associated TFs and lncRNAs.

| Cell state | Total # of conserved associated TFs | TFs supported by other evidences | Total # of conserved associated lncRNAs | lncRNAs supported by other evidences<sup>a</sup> |
| --- | --- | --- | --- | --- |
| ESC | 33 | APEX1, BLM, CARM1, FOXD3, HCFC1, JARID2, LIN28A, MYBBP1A, MYBL2, MYCN, NANOG, POU5F1, POU5F1B, PRDM14, PRDM5, RARG, SALL4, TERF1, TP53, ZFP42, ZNF281, ZSCAN10 | 26 | — |
| Brain | 7 | ARNT1, BCL11A, FEZF2, KCTD1, MAP3K10, NEUROD6 | 34 | — |
| Cerebellum | 17 | BARHL1, BARHL2, LHX1, NEUROD1, ST18 | 58 | TMEM161B-AS1 (ENSG00000247828, blastnOrangutan.Locus_265345) |
| Heart | 12 | ANKRD1, EBF2, GATA4, HAND2, HEYL, MEF2A, MYOCD, NKX2-5, SMYD1, TBX5, TBX20 | 63 | H19 (ENSG00000130600) |
| Liver | 9 | CREB3L3, FOXA3, LPIN2, MLXIPL, NR1H4, NR1I2, NR1I3 | 48 | — |
| Kidney | 12 | GLIS2, HOXA9, HOXD8, HOXD10, PAX8, SIM1 | 69 | LINC00853 (ENSG00000224805), CYP4A22-AS1 (ENSG00000225506) |
| Testis | 41 | BRDT, CREM, FHL5, GTF2A1L, MAK, OVOL1, OVOL2, RFX2, SFMBT1, SOX30, TCFL5, YBX2, ZFY | 477 | TUSC7 (ENSG00000243197) |

This table summarizes the conserved associated TFs and lncRNAs of seven cell states. Conservation definition: human-chimpanzee-bonobo-mouse for TFs; human-chimpanzee-bonobo for lncRNAs. Reported tissue-specific biological functions of some TFs are listed as "other evidence" in the 3rd column of this table. The complete lists of conserved associated TFs and lncRNAs are provided in Supplementary Files 3 and 5, respectively. References are summarized in Supplementary File 4.
<sup>a</sup>Annotated in lncRNADB (http://www.lncrnadb.org).

Although we have identified more conserved associated lncRNAs than TFs (Table 1, Supplementary File 6), fewer lncRNAs have known biological or molecular functions. Hence, our results will provide important new insights into the biological functions of lncRNAs. Three example lncRNAs are summarized in Figure 5A, accompanied with their relative expression levels across seven cell states. We identified H19 as a conserved associated lncRNA of heart, consistent with previous findings that H19 is only expressed in skeletal muscle and heart in human adults (49). We also identified CYP4A22-AS1 (also known as ncRNA-a3) as a conserved associated lncRNA of kidney. Although CYP4A22-AS1 was known to function as an enhancer to stimulate TAL1 gene expression in MCF7 cells (50), its regulatory mechanism in kidney remains unclear. Moreover, NTM-IT was discovered as a conserved associated lncRNA of cerebellum, but its molecular functions are largely unknown. Same as the conserved associated TFs, we also expect those conserved associated lncRNAs to be a valuable resource for future functional studies.

**Figure 5.**
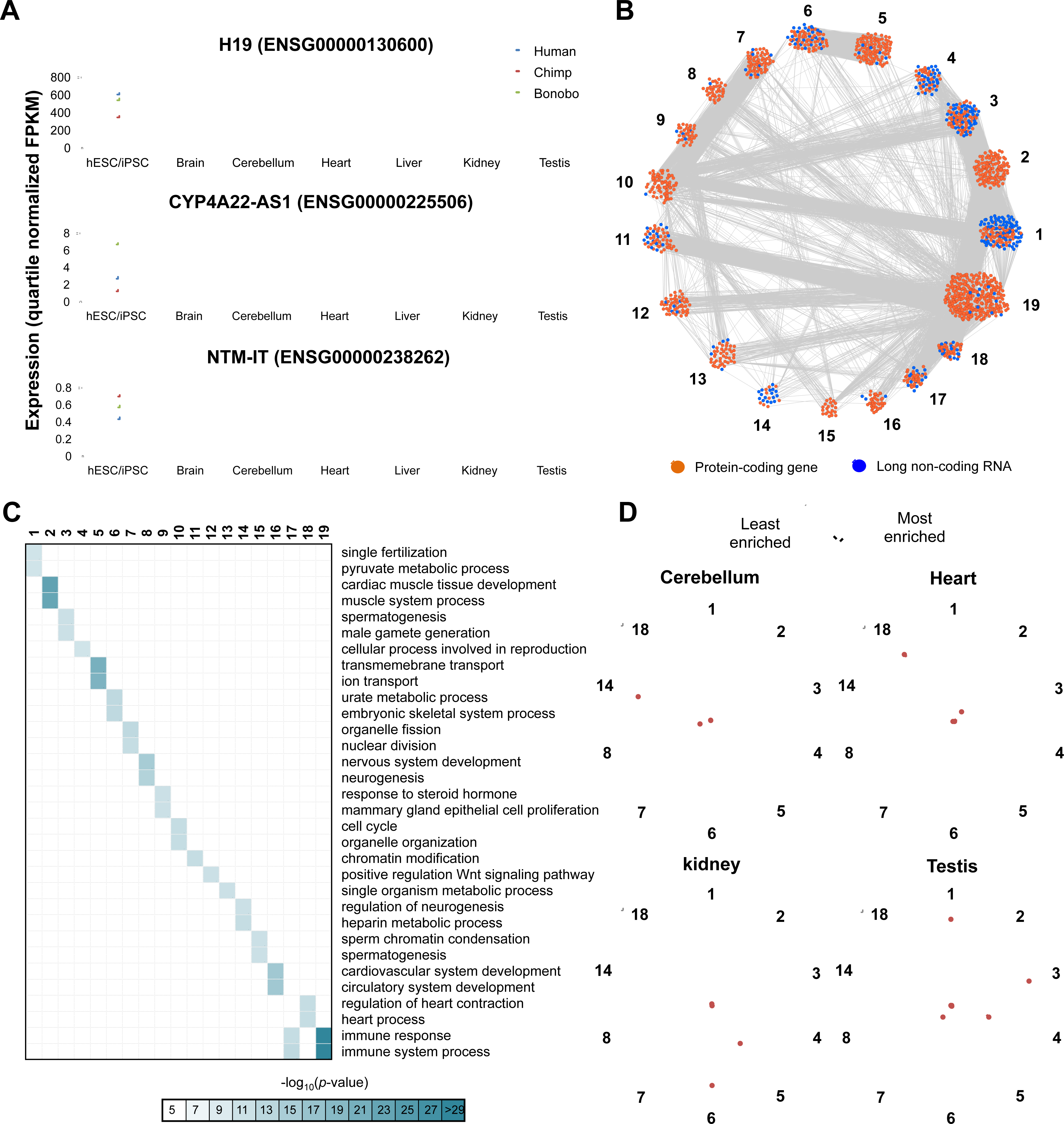
Inferring functions of conserved associated lncRNAs. **(A)** Three examples of conserved associated lncRNAs in heart (H19, ENSG00000130600) (top), kidney (CYP4A22-AS1, ENSG00000225506) (middle) and cerebellum (NTM-IT, ENSG00000238262) (bottom). The expression estimates of the three lncRNAs across seven cell states in human, chimpanzee and bonobo are shown. **(B)** The 19 largest clusters in the conserved co-expression network of protein-coding genes and lncRNAs. Colors of dots distinguish protein-coding genes (orange) and lncRNAs (blue). (**C**) Enriched GO terms (biological processes) of the protein-coding genes in the 19 largest clusters. More significant enrichment scores (defined as –log_10_ transformed *p*-values) are shown in darker colors. (**D**) Rader plots illustrate the extents to which the conserved associated lncRNAs of different cell states are enriched in different clusters. The cell states include cerebellum (top left), heart (top right), kidney (bottom left) and testis (bottom right).

### 3.4 Inferring lncRNA functions using conserved co-expression modules

To infer putative biological functions and regulatory mechanisms of the cell state associated lncRNAs, we constructed a co-expression network of lncRNAs and protein-coding genes. We analyzed a set of 16,354 protein-coding gene families and 13,404 lncRNA families across the three primates. To avoid spurious co-expression interactions, we used evolutionary conservation to determine high-confidence co-expression relationships, which exhibit consistent co-expression patterns across multiple species and thus have selective advantage (51). We evaluated expression correlations for all gene pairs and tested if the cross-species combination of correlations was significantly (*p*-value<0.01) higher than expected by chance (see Methods for details). Finally, the conserved co-expression relationships formed a network with 15,333 nodes (14,369 protein-coding and 964 lncRNAs) and 1,387,285 edges.

We then inferred the potential biological functions of the 964 lncRNAs based on the functions of their co-expressed protein-coding genes. Using the Markov clustering algorithm (52), we identified 19 clusters of highly inter-connected genes (Figure 5B). The protein-coding genes in these clusters are enriched with tissue-specific functions (Figure 5C), such as sperm chromatin condensation, male gamete generation and spermatogenesis in clusters 3 and 15 (testis), nervous system development and neurogenesis in cluster 8 (neural tissues), cardiac muscle tissue development and muscle system process in cluster 2 (heart and skeletal muscle), and immune response and immune system process in cluster 19 (immune cell and tissues). Another interesting finding is that that the conserved associated lncRNAs of various cell states are consistently more enriched with the clusters having the corresponding tissue-specific functions (Figure 5D). For example, the conserved associated lncRNAs in cerebellum are most enriched with the protein-coding genes in cluster 14, whose most enriched biological functions are regulation of neurogenesis and heparin metabolic process. Notably, the clusters with highest lncRNA proportions (clusters 1 and 3) are enriched with spermatogenesis functions, in agreement with the predominant lncRNA specificity in testis (26).

## Discussion

A central goal of developmental biology is to understand the molecular mechanisms underlying the differentiation of ESCs or progenitor cells into various terminally differentiated tissues and cell types. Comparison of gene expression profiles from different cell states would be helpful for deciphering such mechanisms. However, there is in lack of transcriptome-based quantitative metrics to define the similarities or differences between various mammalian cell states, including ESCs, progenitor cells, terminally differentiated tissues and cultured cell lines. Here, we applied a statistical method, TROM, to perform a comprehensive transcriptome mapping using RNA-seq data from diverse tissues and cell types within and across four mammalian species. We identified associated protein-coding genes as good cell state markers, which capture transcriptome characteristics and lead to reasonable correspondence maps of cell states both with-species and between-species. We also identified associated lncRNAs, which are found as good cell state markers as well and even more cell state specific than protein coding genes. In addition, we constructed a cell differentiation tree based on the transcriptome similarity of various human cell states. Furthermore, we compared the transcriptomes of various cell states across multiple mammalian species using TROM, and identified conserved associated protein-coding genes and lncRNAs. Finally, we investigated the evolutionary signatures of those conserved associated genes of various cell states, and confirmed known and discovered potential biological functions for them. In addition to uncovering several novel insights into mammalian transcriptomes, this study also provides a useful resource of conserved associated protein-coding genes and lncRNAs, which characterize transcriptomes of various cell states and enable researchers to explore new hypotheses in developmental biology and evolutionary biology.

We realize that our current cell state mapping results are limited in two aspects. First, our tissue samples are heterogeneous populations of cells, such as the brain tissues containing multiple types of neurons. Because of that, our mapping results cannot well detect neuronal subtypes, though we partially resolved this issue by using lncRNAs or TFs, which exhibit higher tissue-specificity expression patterns than protein-coding genes. In addition, we found that some samples have obvious within-tissue variation. For example, some liver samples from chimpanzee have a stronger mapping to spleen and kidney than to liver tissues in mouse. This result is probably due to the fact that liver tissues include fibroblasts, vascular tissues and many other cell types in addition to hepatocytes. We anticipate that this heterogeneity issue could be resolved with the availability of single-cell RNA-seq data. Despite this limit, our mapping strategy performs well in assessing transcriptome similarities across a wide range of mammalian tissues and cell lines, as demonstrated by our results.

Second, lncRNA expression estimates are not as accurate as the protein-coding gene expression estimates, because lncRNAs have much lower expression levels than those of protein-coding genes. Many previous studies have revealed that the median expression level of lncRNAs is only about a tenth of that of mRNAs (32, 53-58). This makes accurate estimation of lncRNA expression a generally difficult task. Our curated RNA-seq data sets were generated over a span of more than five years, resulting in a great variety in their sequencing depths and read lengths. Notably, the data we used from early studies are short read RNA-seq (<50 base pair single-end reads) of moderate sequencing depths (∼10-20 million reads). Advances in RNA-seq library preparation and sequencing technologies now enable the generation of hundreds of millions of paired-end reads with longer than 100 base pairs in length. Greater sequencing depths and increased read lengths allow more accurate abundance assessment of lowly expression genes (59), and thus could improve our cell stage mapping results.

Past years have seen great progress in studying cell-state transitions. For example, fibroblasts (60) and neural progenitor cells (61) have been reprogrammed to pluripotent stem cells, which can again differentiate into specific lineages through defined growth conditions and/or gene expression perturbation (62). Some previous studies revealed that knocking down key regulatory factors in the initial cell state could aid the cell state conversion (6, 63). However, those studies were based on microarrays and not comprehensive. Our study for the first time provides the repertoires of various cell states’ associated protein-coding and non-coding genes identified based on RNA-seq technologies. We anticipate that these genes are valuable candidates for further functional studies to improve the efficiency and fidelity of cell-state conversion and reprogramming (48).

In addition to the increasingly accumulation of RNA-seq data, a large amount of microarray data sets have been produced and deposited during the past decade. To this end, our cell state mapping approach allows us to integrate RNA-seq and microarray data for studying more cell states in more species. Our method can also be extended to incorporate additional epigenomic data types, such as DNA methylation, histone modification, DNA accessibility and genome-wide transcription factor binding data. Some recent studies demonstrated that cell and tissue lineages could be well clustered according to their DNA accessibility (39) and histone modification (64) patterns. Ongoing efforts to further the knowledge of cell-state associated regulatory genes will greatly advance our understanding of ESC state maintenance, as well as our ability in cell reprogramming and directed differentiation.

## Methods

### RNA-seq data collection and processing

We compiled a data resource of 307 publicly available poly(A) RNA-seq data sets, which were profiled from four mammalian species: *Homo sapiens* (human), *Mus musculus* (mouse), *Pan troglodytes* (chimpanzee), and *Pan paniscus* (bonobo). Our data resource contained ∼19 billion sequencing reads from 307 biological samples, of which seven cell and tissue types (brain, cerebellum, heart, kidney, liver, testis and ESC or iPSC) existed in all four species. In addition, our data resource covered far more cell and tissue types in human (totally 40 tissues and cell types) and mouse (totally 18 tissues and cell types). All RNA-seq data sets were generated from Illumina GAII or HiSeq2000/HiSeq2500 systems. We filtered out low quality reads in each RNA-seq data set using PRINSEQ (65).

We then aligned the RNA-seq data to the mammalian genomes using Tophat v2.0.10 (66, 67). Three mammalian genomes were obtained from Ensembl (*Homo sapiens* hg19, *Mus musculus* mm10, and *Pan troglodytes* panTro4). Since the genome of bonobo (*Pan paniscus*) was not available in Ensembl, we used the chimpanzee genome (panTro4) as the reference genome for bonobo in our analysis, given an observed high (>95%) similarity between bonobo and chimpanzee genomes (68). The detailed mapping results were in Supplementary File 1.

### Gene expression estimation

We constructed mammalian reference genome annotations by integrating published annotations from multiple resources. For protein-coding genes, the annotations were from human Gencode v19, mouse Gencode M2, and chimpanzee/bonobo Ensembl_CHIMP2.1.4.73 (69, 70). For lncRNAs, the annotations were from a recent publication (26). We then used Cufflinks (30, 66) (v2.1.1, supplied with reference annotation, i.e., using “-G” option) to measure the expression levels for both protein-coding genes and lncRNAs in the four mammalian species.

Since RNA-seq data sets from public data repositories were generated in different studies by different laboratories, we used a unified procedure to evaluate the gene expression estimates (in FPKM (fragments per kilobase of transcript per million mapped reads) units) (30, 66). By examining the reproducibility of the FPKM values between replicates within the same tissue or cell type, we found high correlation coefficients between all replicate pairs. LncRNAs exhibited much lower expression levels and higher tissue- or cell-type specificity than protein-coding genes, consistent with previous results (32, 55, 71, 72). These results show that our data resource is a proper basis for further analyses.

### Orthologous gene families and TF annotations

We obtained orthologous families of protein-coding genes from TreeFam v9 (73), a database of tree-based orthology predictions, and orthologous families of lncRNAs from a recent publication (26). The orthologous families are summarized in Supplemental File 2.

For TF analysis in our studies, we downloaded TF annotations of human, mouse, chimpanzee and bonobo from AnimalTFDB 2.0 (74), which provides TF annotations for more than 50 animal genomes.

### Identification of cell state associated genes by TROM

One cell differentiation state (i.e., a tissue or cell type, such as ESC, liver, testis, etc.) can be characterized by its “associated genes”, which we defined as the relatively highly expressed genes in cell state as compared to other cell states. We first calculated relative expression estimate (i.e. *Z*-score) across cell states for each gene. Next, we normalized the *Z*-score distributions of different cell states within every species using quartile normalization (75). Then for every cell state, we selected its associated genes whose FPKMs is above a threshold and normalized *Z*-scores are in a top percentile. The FPKM threshold is 1.0 for protein-coding genes and 0 for lncRNAs, and the *Z*-score percentile is top 5%. These two criteria guarantee that in the given cell state, the expression levels of the associated genes are distinguishable from background noise and are also higher than in some other cell states.

We then used the TROM algorithm to map two mammalian cell states by their transcriptome similarity (28). In the within-species cell state mapping, for every two cell states, we tested the significance of the number of their common associated genes using a hypergeometric test, whose null hypothesis is that the two cell states’ associated genes are independent samples from the gene population of the species. If significant, two cell states were called mapped. In the between-species cell state mapping, for every two cell states from different speices, we tested the significance of the number of ortholog pairs between their associated genes using an approximate hypergeometric test. If the test was significant, two cell states were called mapped.

### Construction of cell differentiation tree

Based on the associated genes we identified, we constructed cell differentiation trees of different cell differentiation states in human. We selected 27 representative cell states of three categories: (i) ESCs and iPSCs, the precursors of all the other tissues and cell types, (ii) germ layers and other progenitor cell types, and (iii) differentiated tissue types (e.g., liver, kidney, brain, etc.).

We constructed the cell differentiation trees using associated protein-coding genes and TFs respectively following a previously proposed method (76). First, we calculated the Jaccard distance between pairwise cell states and applied hierarchical clustering with average linkage to cluster the cell states. Second, we used the Hungarian algorithm (77) to place the progenitor cell types to the best-fitting nodes of the constructed clustering tree.

### Enrichment analysis of Gene Ontology and biological pathways

For Gene Ontology (GO) analysis, we used topGO (78) to estimate the enrichment of Biological Process (BP) terms for different cell states based on their associated genes. We calculated the significance of GO term enrichment in every cell state using a hypergeometric test. The top three most enriched GO terms in every cell state were displayed in Figure 2C and Supplementary Figure 9-11. For biological pathway analysis, we used KEGGREST (79) to calculate the enrichment of biological pathways, and displayed the results in a similar fashion as in the GO analysis.

### Identification of conserved cell state associated TFs and lncRNAs

To identify conserved cell state associated TFs, we used the orthologous families of the TFs in human, chimpanzee, bonobo and mouse. As an example, suppose TF *X* is a liver-associated TF in human. If its orthologous genes in the three other species are also liver-associated TFs, it is defined as a conserved associated TF of liver. Overall, we identified conserved associated TFs in seven cell states: ESC, brain, cerebellum, heart, liver, kidney and testis.

We used the same approach to identify conserved cell state associated lncRNAs, except that the conservation is only based on the three primates, because little is known about the orthology of lncRNAs between primates and mouse. We identified conserved associated lncRNAs in the same seven cell states.

### Construction of conserved co-expression network of protein-coding genes and lncRNAs

We applied a previously published method (51) to reconstructing a conserved co-expression network of protein coding genes and lncRNAs across human, chimpanzee and bonobo. We did not include mouse because there are few known conserved lncRNAs between mouse and primates. We used the well-established homologous families for both protein coding genes (from TreeFam v9 (73)) and lncRNAs (from (26)) across the three species. We then computed the Pearson Correlation Coefficients of expression patterns. Given two homologous families, we examined whether the combination of correlation coefficients from the three species was significantly (*p*-value<0.01) higher or lower than expected by chance. We carried out statistical tests by comparing the observed ranks of the correlation coefficients with random *n*-dimensional order statistics. We computed correlations only for those protein-coding genes and lncRNAs expressed in at least two cell states of each species. Genes were defined as expressed in a cell state if their expression estimates were in the top 90% of that cell state. We allowed for negative correlations in the co-expression network, which is visualized by Cytoscape (80) in Figure 5B.

### GO enrichment in the co-expression network

We identified clusters of highly inter-connected gene sets in the co-expression network via the Markov Cluster (MCL) algorithm (52). Each cluster contains both protein-coding genes and lncRNAs. Then, we detected enriched GO (Biological processes) terms in the protein-coding genes of each cluster. We also calculated the enrichment significance of the conserved associated lncRNAs of several cell states in each cluster using hypergeometric tests. The results were visualized using radar plots in Figure 5D.

## Authors’ contributions

JJL and ZJL conceived and designed the project; YY and YTY performed all the analyses; JY helped in the co-expression network analysis; YY and YTY prepared the figures and tables; JJL, ZJL, XS, YTY and YY wrote the manuscript.

## Acknowledgements

YY, YTY, JY and ZJL were supported by the National Key Basic Research Program of China [grant number 2012CB316503]; the National High-Tech Research and Development Program of China [grant number 2014AA021103]; the National Natural Science Foundation of China [grant numbers 31271402]; Tsinghua University Initiative Scientific Research Program [2014z21045]. JJL was supported by the startup funding provided by the Department of Statistics at UCLA. This work was also supported by Computing Platform of National Protein Facilities (Tsinghua University). We would also like to thank Dr. Mark Biggin for helpful discussions; other members from Lu Lab and Li Lab for their insightful comments and suggestions.

